# Exploring sex-related Biases in Deep Learning Models for Motor Imagery Brain-Computer Interfaces

**DOI:** 10.64898/2026.03.05.709808

**Authors:** Bruno J. Zorzet, Victoria Peterson, Diego H. Milone, Rodrigo Echeveste

## Abstract

Motor imagery (MI) brain-computer interfaces (BCIs) are promising technologies for neurorehabilitation. In this context, deep learning (DL) models are increasingly being used to decode the mental imagination of movement. However, countless studies across multiple domains have shown that DL models are susceptible to bias, which can lead to disparate performance across subpopulations in terms of protected attributes, such as sex. The reported presence of sex-related information in electroencephalography (EEG) signals, widely used for MI-BCI, further raises warnings in this regard. For this reason, we conducted an in-depth analysis of the performance of DL in terms of the sex and other potential confounding factors. While an initial basic stratified analysis in terms of sex showed differences in favor of the female population, further analysis revealed that performance disparities were actually primarily driven by the discriminability of EEG patterns themselves, and not by the DL model. Moreover, DL models improve overall performance as well as per-group performance, particularly helping subjects with less discriminable EEG patterns. Our work highlights the benefits of DL methods for MI-BCI as well as the need for careful analysis when it comes to bias assessment in complex settings where multiple variables interact. We argue that in-depth studies of model behavior beyond standard performance metrics, should become widespread in the community in order to ensure the development and later deployment of fair BCI systems.

## 1 Introduction

Brain-Computer Interfaces (BCIs) are systems that allow direct communication between the brain and external devices. By means of a BCI, brain activity is captured and processed to be converted into controlling commands [1]. Different neuroimaging and signal acquisition technologies are used to register brain activity. Among them, electroencephalography (EEG) is widely used in the context of BCI, since it provides good temporal resolution while being more accessible and portable than other brain recording technologies [2]. In EEG-based BCI systems, the most common paradigms used to convert human intentions into commands are P300 evoked potentials [3], steady state visual evoked potentials [4], and motor imagery (MI) associated potentials [5]. These paradigms present some challenges due to the intrinsic characteristics of EEG signals, such as low signal-to-noise ratio (SNR) [6], non-stationarity [7], and intersubject variability. In this regard, MI signals are particularly difficult to decode since they are produced by the self-regulation of brain activity at different frequency bands.

Considering these challenges, machine learning (ML) methods become good options for decoding complex information from brain data. In particular, deep learning (DL) models are able to capture multiple representational levels in the data, and are highly effective for solving countless tasks as demonstrated in other fields such as medical diagnosis [8] and natural language processing [9]. Due to the larger number of accessible EEG datasets and the development of artificial neural networks with reliable performance, the BCI community is increasingly devoting efforts to using DL approaches for decoding MI EEG signals. The hope is that automatic feature learning, provided by end-to-end trained models, could ultimately lead to more robust classification also in the BCI field. The general superiority of DL over traditional methods in BCI-MI is a well-established fact. Then, the core focus and novelty of this work is to investigate a critical unanswered question: whether the use of DL models in MI-BCI introduces or amplifies demographic biases, specifically related to biological sex, within the context of motor imagery.

For a number of years, DL models have shown to be susceptible to biases with respect to certain sub-populations, leading to disparate performance across different groups defined by protected attributes such as age, biological sex, ethnicity and socio-economic level [10]. Due to their large amount of trainable parameters, DL models are highly powerful tools to learn inner representations of the data. Interestingly, this same complexity makes them prone to misleadingly amplify irrelevant information. This issue gave rise to the field of algorithmic fairness, which aims to detect (and ideally mitigate) potential sources of harmful bias. This is crucial in fields such as healthcare, where such disparities can be considered to go against bioethical principles of justice, autonomy and nonmaleficence [11]. Indeed, in domains such as medical image computing, it was demonstrated that DL approaches can improve the performance of the model for one subpopulation to the detriment of others [12, 13].

When it comes to MI-BCIs as well, the assessment of model performance should take into account the existence of potential biases with respect to demographic attributes of users. The amplification of such disparities could arise not only from the model itself but also from the training data, the training process, and the design choices made during model development [13]. For instance, if the training data is not representative or is imbalanced in terms of demographic factors, this could lead to biased learning outcomes. Similarly, the way in which the model is trained or optimized might exacerbate these biases. The assumptions of developers, choices in model architecture, and the features selected for training could also influence how these disparities are reflected in the final model.

Multiple studies in the past have evidenced that protected attributes can themselves be decoded from the data, potentially resulting in biased machine learning systems. Issa et al. managed to classify the sex and age of the participants from EEG signals with a simple preprocessing wavelet transform, achieving accuracies of 0.89 and 0.96 for sex and age predictions, respectively [14]. Kaushik et al. reached accuracies of 0.93 and 0.97 in predictions of age and sex from EEG signals using a DL model based on a combination of Bidirectional Long-Short Term Memory (BLSTM) and Long-Short Term Memory (LSTM) layers, demonstrating the risk of DL models finding subtle statistical regularities [15]. Kaur et al. achieved accuracies of 0.83 and 0.97 in age and sex predictions using a discrete wavelet transform on EEG signals. They also explained that beta and theta waves contribute to the classifications of age, while delta rhythms contribute to the classification of sex [16]. Wang and Hu reached accuracies of 0.99 and an area under the receiver operating characteristic curve (AUC-ROC) of 0.99 for sex recognition using only a random forest and logistic regression on EEG signals during resting state [17]. Finally, Van Putten et al. showed that deep neural network models can achieve accuracy over 0.80 in sex classification from EEG signals [18].

The fact that it is possible to identify protected attributes such as age and biological sex from EEG signals is certainly a cause for concern for the fairness community, but it does not necessarily mean that systems will be biased. Furthermore, most work so far has focused on resting state EEG signals, rather than evoked signals such as those core to MI-BCI. Only recently, biases in classification tasks involving intransitive (i.e., meaningful gestures that do not include the use of objects), transitive (i.e., actions involving an object), and tool-mediated (i.e., actions involving a tool to interact with an object) movements of the upper limbs were studied. Catrambone et al., for instance, found differences in the performance of a k-NN classifier predicting movements of upper limbs between females and males. They observed a significant difference in accuracies favoring female subjects [19]. However, to our knowledge there are no studies that evaluate the presence of biases in DL models for brain decoding in the context of MI-BCI tasks.

Analyzing whether the performance of DL models in MI tasks may be influenced by sex, age or other protected attributes is essential to ensure the aforementioned bioethical principles. This is crucial in applications like MI-BCI for rehabilitation [20], where certain sub-groups of patients recovering from a stroke may be disproportionately affected by biases in the model. Nevertheless, in MI-BCI users must self-regulate their brain activity, which is influenced by their emotions, experiences and feelings. Consequently, some individuals find MI tasks easier to perform than others [21]. In other words, from the machine learning viewpoint, data quality strongly depends on MI-BCI user skill. Therefore, in order to distinguish potentially confounding sources of bias it is crucial to quantify MI performance of individuals, independently of the ML model used to decode such brain data [22]. In this way, the ability of the user to self-regulate their brain activity can be understood as another attribute of the sample population.

Here, we focused on across-subject paradigms for MI-BCI, not only evaluating disparities in performance between males and females but also carrying out statistical analyses to detect these potential confounders. Our findings suggest that while DL models can amplify differences in performance across subpopulations, these differences are predominantly driven by the intrinsic user capability to generate discriminative MI related patterns, rather than by demographic attributes such as biological sex. Indeed, we show how spurious correlations between sex and signal discriminability may lead to misleading conclusions if one were to conduct a naive analysis of performance across groups. This study represents a step forward in understanding the variables that can potentially influence the performance of MI-BCI decoding algorithms. Moreover, it underscores the importance of developing fair DL models that not only achieve high performance but also mitigate potential inequalities. All code used in this study is publicly available at: GitHub.

## 2 Materials and Methods

### 2.1 Dataset and Preprocessing

Two MI-BCI datasets were used in this study. Both of them comprise the mental imagination of left vs. right hand movement. These datasets were selected based upon: number of participants, open accessibility, and metadata information availability (sex and age of participants).

#### Lee 2019[24]

EEG signals were acquired at a sampling rate of 1000 Hz using 62 Ag/AgCl electrodes according to the international system 10-05. The channels were nasion-referenced and grounded to electrode AFz. This dataset includes data from 54 subjects (mean age = 24.2 ± 3.04 years, 23 females). The experiment consisted of two sessions of MI tasks, at which a training and a testing phase were conducted. Only data from the training phase, which included label information, was used in this study. Each participant performed 100 trials of left vs right MI (50 trials per class) at each training phase and session. The MI-BCI paradigm followed established protocol standards in which after a visual cue, 4 s were given to the participant to perform MI. No feedback was provided to the users.

#### Cho 2017[23]

EEG signals were acquired at a sampling rate of 512 Hz. 64 Ag/AgCl electrodes according to the international 10-05 system were used. The dataset has 52 participants (mean age = 24.8 ± 3.86 years, 19 females). Each participant performed five or six runs of 20 trials of hand movement imagination tasks. Authors used a standardized MI-BCI protocol, where after a visual cue participants had 3 s to perform the mental imagination, without any followed type of feedback over trials.

Further demographic details for both datasets are provided to characterize the participants. The age distribution of the subjects across the Lee 2019 and Cho 2017 datasets is illustrated in Figure S3. Furthermore, the previous BCI experience of the participants is summarized in Table S8.

Data was pre-processed following a standard pipeline for MI-BCI decoding. First, a zero-phase band-pass filter between 0.5 - 40 Hz was applied over raw data, followed by a Notch filter at 50 Hz. Given that each dataset used different positions for the reference electrode, in both datasets, channels were re-referenced to Fz. In addition, signals were downsampled to 128 Hz. Also, EEG windows were cropped between 0.5 to 2.5 s after the MI cue. To simplify the scenario, only channels C3, C4 and Cz were selected. Once this pre-process was applied, each dataset consisted of EEG trials from 3 channels with 256 time samples.

## 2.2 Model

Due to its wide acceptance and application for EEG decoding [36, 37, 38], here we adopted EEGNet, a convolutional neural network for EEG-based BCIs, as a reference DL network architecture. This DL architecture receives as inputs raw EEG data of dimension (*C, T*), where *C* is the number of channels and *T* is the number of time points in each trial. It follows a traditional MI detection pipeline implemented by a series of convolutions, starting from a temporal convolution, aiming at filters the signals of each channels, which can be viewed as a temporal bank. To aggregate information from all channels, a depthwise convolution follows the filter-bank, mimicking the well-known spatial filtering techniques, as the common spatial patterns (CSP) method [30]. Lastly, the model performs a separable convolution layer and pointwise convolution, to aggregate information across feature maps. The extracted features from all previous layers are mapped to the output classes (here left hand and right hand MI) by the final fully connected layer.

The analysis we present in the following for the EEGNet was however extensively validated on multiple other models (See Supplementary Material). In particular, we included: Deep4Net [26], ShallowFBCSPNet[26], HybridNet [26], FBCNet [27], CTNet [28], ATCNet [29] and the traditional CSP+LDA method.

## 2.3 Experimental setup and training process

We trained the EEGNet for binary MI classification using an across-subject scheme. For each dataset we applied a leave-one-subject-out (LOSO) evaluation scheme as described below. Each dataset can be defined as a set 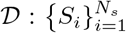, where *S*_*i*_ = (**X**_*i*_, **y**_*i*_) represents the data of every subject, that is, the EEG trials during the MI task **X**_*i*_, and the corresponding labels, **y**_*i*_, and *N*_*s*_ stands for the number of subjects in a dataset. The tensor **X**_*i*_ has dimensions (*N*_*i*_, *C, T*), where *N*_*i*_ is the number of trials for subject *i*, and *C* and *T* represent the number of channels and time points, respectively. The labels vector is defined as 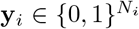 where 0 is for left hand and 1 for right hand. Additionally, the metadata information is represented by 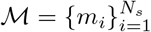, where *m*_*i*_ ∈ {0, 1}, with 0 indicating females and 1 indicating males.

To ensure balanced data during training and evaluation, for each subject *S*_*i*_ the following process was repeated for *N*_*r*_ = 20 replicates:

1. Conform the subset for training and validation by including the remaining subjects in the dataset. Mathematically, 𝒟_*−i*_ = {*S*_*j*_ | ∀*j* ≠ *i*} and ℳ_*−i*_ = {*m*_*j*_ | ∀*j* ≠ *i*}. The dataset 𝒟_*−i*_ and metadata ℳ_*−i*_ is then randomly balanced with respect to biological sex using a function which ensures that both training and validation subsets have a balanced representation of male and female participants. As outputs, a subset of ignored participants (𝒟_*ign*_ and ℳ_*ign*_) as well as a balanced subset (𝒟_*bal*_ and ℳ_*bal*_) are obtained.
2. From the balanced subset, data from *N*_*sv*_ = 4 subjects (two male and two female) are randomly selected to construct the validation set, hence preserving the balance in terms of biological sex in both training and validation sets. As customary, the validation set is used to monitor overfitting during training (see details of datasets creation in Algorithm 1).
3. For each training and validation subset, the model is trained with *N*_*sd*_ = 5 different initialization seeds, so as to be able to desegregate the variance with respect to data from the variance with respect to model training. The final model is evaluated using data from the leave-out subject *S*_*i*_.

As a result of this process, *N*_*r*_ = 20 partitions of training and validation, with *N*_*sd*_ = 5 different random initializations of the models were obtained, totaling *N*_*m*_ = 100 models for each leave-out subject. This method guarantees that every model of every test subject, independently of the biological sex, is trained by the same number of subjects, and all data is balanced by sex.

The explicit balancing of subjects by sex within the LOSO training partitions is a methodological choice designed to prevent disparities arising purely from data imbalance. By enforcing balance across all *N*_*r*_ replicates, we ensure that the model is trained with an equal opportunity to learn from both male and female participants, thereby allowing us to isolate and study the performance disparities caused by subtle factors, such as feature heterogeneity. For a separate study that analyzes the impact of data imbalance on biases see Supplementary Material, Section S4.2.

Cross-entropy was used as the loss function for binary classification, and Adam [31] was used as an optimizer with its default settings, and a learning rate of 0.0001. To account for a traditional machine learning pipeline from which compare performance, the gold-standard CSP followed by a linear discriminant analysis (LDA) was also used for MI decoding. A pair of CSP components were extracted. Both models were trained and evaluated using the same data partitions

### Algorithm 1

Create balanced data split by biological sex

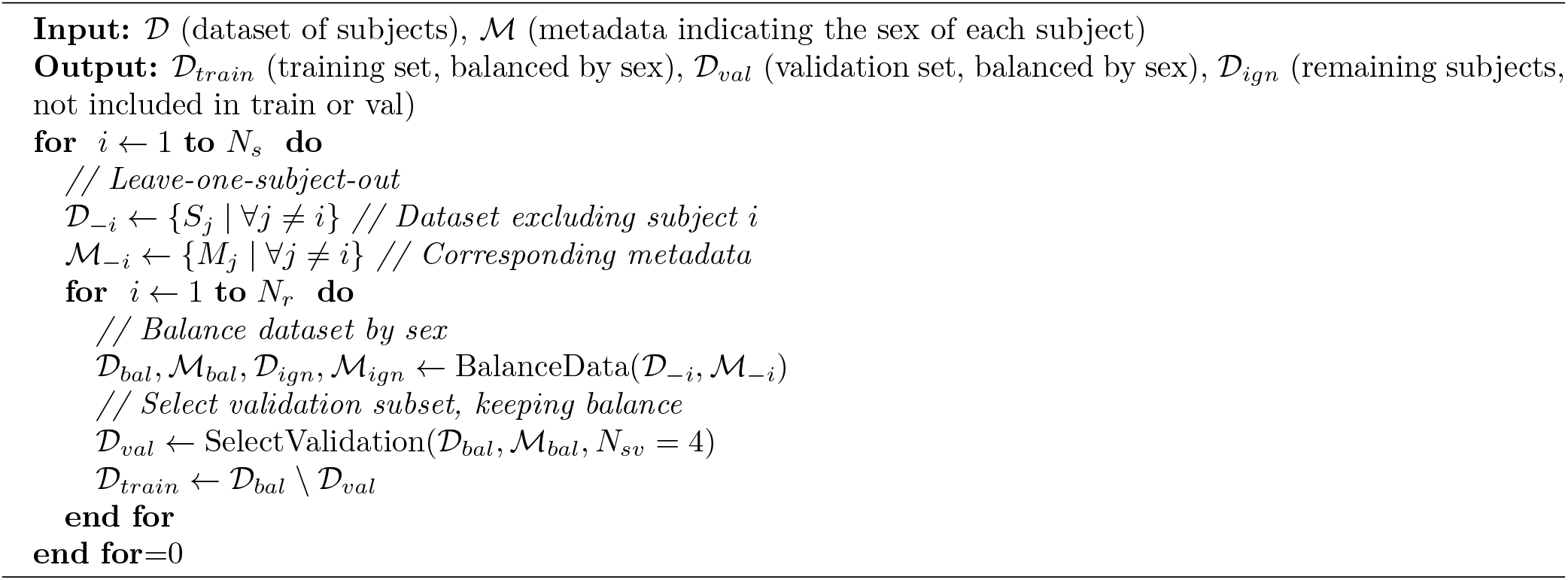

## 2.4 Performance metrics and statistical analysis

In what follows we quantify the performance of the models for each subject, both in terms of accuracy and area under the ROC curve (AUC). It is worth mentioning that these metrics reflect a combination of the effectiveness of the models themselves and the skill level of the BCI users to modulate their brain signals. Therefore, it is crucial to be mindful of both contributions when interpreting performance metrics.

In order to have a separate and independent measure of the modulatory abilities of subjects, we compute the class distinctiveness of the signals. Unlike accuracy or AUC, class distinctiveness is a metric based purely on the signal, computed directly over the raw EEG input, and hence it is independent of the deep learning model. It quantifies the distance between the distributions of right hand and left hand MI trials in a Riemannian space, thus acting as a direct proxy for the difficulty of the classification task. For every subject *S*_*i*_, we used the tensors **X**_*i*_ of *N*_*i*_ trials for subject *i, C* channels and *T* samples point to compute the spatial covariance 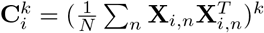 matrix, defined as for every class *k*.

With this matrix, we can compute the class distinctiveness as:

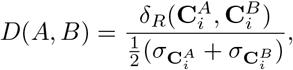

where *δ*_*R*_ is the Riemann distance between the spatial covariance matrix of each class and 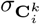 is the class dispersion of class *k*. Each spatial covariance matrix was calculated with the already preprocessed data as explained above, considering only the same subset of channels used to train the models. Class distinctiveness was calculated for every train and validation set of signals of each test subject. This allowed us to separate the contribution of class distinctiveness of the EEG patterns in the training and validation data, from the impact determined by the distinctiveness in the test data.

As LOSO was used to evaluate classifier performance, there is high overlap in the subjects in the training sets corresponding to different test subjects. This overlap can compromise the independence between test performance metrics points, posing challenges for statistical tests designed to assess the significance of model performance differences. Traditional tests, such as t-tests, ANOVA or ANCOVA, assume independence of observations; deviations of this assumption reduce statistical power. In addition, we aimed to understand the sources of differences in performance metrics in relation to covariates such as sex, age, and class distinctiveness. To achieve this, we employed mixed-effects models [32], which account for the non-independence of data by incorporating fixed effects for the covariates of interest and modeling subject-level variability as a random effect. This approach provides a more robust framework for quantifying the contribution of each covariate to the distribution of performance metrics and offers deeper insights into their impact on model performance.

## 3 Results

Across both datasets, we observed that the performance of the DL model (exemplified here by the EEGNet) is higher than CSP+LDA, both globally and when segregated by sex. Additionally, a consistent trend of higher accuracy scores for female participants compared to male participants was evident (cf. Male vs. Female in Table 1 and in Table S1). When comparing the differences between the two models, we observed that the performance gap between sexes for CSP+LDA is relatively small, ranging from 1% to 4%. However, this occurs in a context where the overall performance metrics are notably poor, limiting the utility of the model. In contrast, DL models exhibit larger performance gaps, ranging from 5% to 7%, but also achieve significantly higher overall performance metrics for both groups. This trade-off highlights a critical point: while DL models may introduce greater disparities across subpopulations, they also provide substantial improvements in classification accuracy, making them more effective in practical applications despite potential bias. Similar trends were observed in AUC metrics (see Table S2), where the differences between models and subpopulations were even more pronounced.

**Table 1:** Accuracy performance comparison of different models (EEGNet and CSP+LDA), showing global accuracy and segregated by sex. A Mann-Whitney U test was applied to assess differences between female and male subjects, and the resulting p-values were adjusted using the Bonferroni correction due to the number of comparisons (*n* = 8).

| Datasets | Models | Global Accuracy | Females | Males | p-values |
| --- | --- | --- | --- | --- | --- |
| Cho 2017 | EEGNet | $0.71 \pm 0.13$ | $0.75 \pm 0.13$ | $0.68 \pm 0.12$ | 0.44 |
| | CSP+LDA | $0.59 \pm 0.15$ | $0.62 \pm 0.14$ | $0.58 \pm 0.15$ | 0.99 |
| Lee 2019 Session 1 | EEGNet | $0.71 \pm 0.12$ | $0.74 \pm 0.11$ | $0.69 \pm 0.12$ | 1.00 |
| | CSP+LDA | $0.59 \pm 0.08$ | $0.59 \pm 0.09$ | $0.58 \pm 0.07$ | 1.00 |
| Lee 2019 Session 2 | EEGNet | $0.71 \pm 0.12$ | $0.74 \pm 0.11$ | $0.69 \pm 0.12$ | 1.00 |
| | CSP+LDA | $0.55 \pm 0.08$ | $0.56 \pm 0.08$ | $0.54 \pm 0.08$ | 1.00 |

To better understand the performance gaps, we computed the logarithm of class distinctiveness for each test subject, grouped by sex, as summarized in Table 2. This transformation addresses the highly skewed distribution of class distinctiveness (see Figure S1 in Supplementary Material), allowing for a better comparison, not dominated by the tails of the distribution.

**Table 2:** Average class distinctiveness segregated by sex. A Mann-Whitney U test was performed to assess whether differences between sexes exist.

| Dataset | Female | Male | p-value |
| --- | --- | --- | --- |
| Cho 2017 | $-2.45 \pm 1.66$ | $-3.16 \pm 1.74$ | 0.11 |
| Lee 2019 Session 1 | $-2.53 \pm 0.90$ | $-2.83 \pm 0.82$ | 0.53 |
| Lee 2019 Session 2 | $-2.68 \pm 1.25$ | $-2.60 \pm 1.11$ | 0.69 |

Statistically, there are no differences between female and male populations. In the Cho 2017 dataset the female population as whole has a higher class distinctiveness, in accordance with their superior performance in terms of accuracy and AUC, but in Lee 2019 this is not the case. These results showed the complex relationship between MI capabilities and model decoding performances, highlighting the need for a more in depth analysis.

To examine the influence of class distinctiveness and sex on performance metrics in further detail, we computed correlations between performance metrics and class distinctiveness. The analysis was performed separately for each dataset and session, with results segregated by sex. Additionally, we conducted partial correlation analysis using sex as a covariate to isolate the relationship between performance and class distinctiveness, minimizing potential confounding effects of sex. In Figure 1, scatterplots depict the average accuracy of each test subject versus the class distinctiveness for DL models, CSP+LDA, and their difference. For the Cho 2017 dataset (first row in Figure 1), accuracy strongly correlated with the class distinctiveness, with *r* = 0.86 for CSP+LDA and *r* = 0.83 for EEGNet. When segregated by sex, males exhibited stronger correlations than females for both CSP+LDA (*r* = 0.87 vs. *r* = 0.85) and EEGNet (*r* = 0.85 vs. *r* = 0.79).

**Figure 1.**
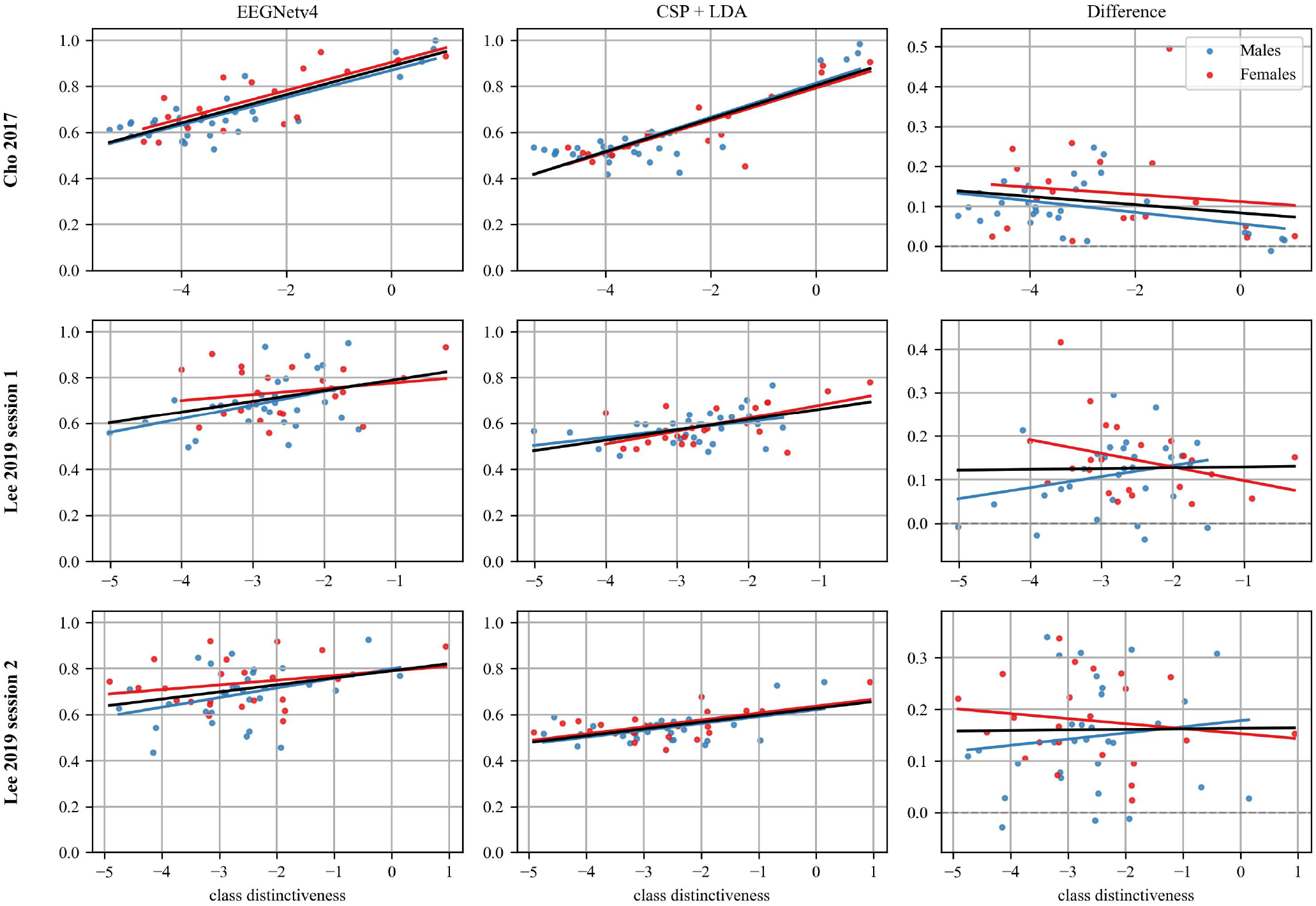
Scatterplots of accuracy vs. class distinctiveness for both databases. Each row corresponds to a specific database, and the columns represent the accuracy performance of the DL (EEGNet) model, CSP+LDA, and their differences. Each dot represents the average performance across all models trained to test for a single subject. Data points for males are shown in blue, while those for females are in red. Straight lines indicate regression trends: black lines represent regression across all data (not segregated by sex), and colored lines correspond to sex-specific regressions.

For the Lee 2019 dataset (Figure 1 second and third row), correlations were weaker compared to Cho 2017. In Session 1, CSP+LDA achieved *r* = 0.52, while EEGNet showed *r* = 0.36. Within subgroups, females had stronger correlations for CSP+LDA (*r* = 0.60) but weaker correlations for EEGNet (*r* = 0.22) compared to males (CSP+LDA: *r* = 0.42; EEGNet: *r* = 0.42). In Session 2, males consistently showed stronger correlations than females for EEGNet model. Regarding AUC, correlations were generally higher, with dataset-specific variations, see Figure S2 and Table S4.

From Figure 1, we observe that for CSP+LDA the regression lines for males and females are similar, which aligns with the small differences between their mean performance in each dataset. However, when comparing the differences between CSP+LDA and EEGNet metrics, the improvement in accuracy is consistently higher for females than for males. In the Cho 2017 dataset, the regression line for females is consistently higher than that for males across the entire range of data. In the Lee 2019 dataset, the regression line for males surpasses that for females at a class distinctiveness around − 2. Similarly, in Session 2, this crossover occurs at approximately − 1. However, it is worth noting that the scatter plot points for females are often positioned below their regression line. This is attributed to the smaller number of data points in that range, which could cause the regression curve to overfit their population. Further details are available in Table S3 and Table S4.

In order to isolate the effects of class distinctiveness on performance metrics, we compute partial correlations to assess the relationship between two variables while removing the effects of covariates (such as sex) [33]. The results showed that class distinctiveness remained a consistent factor influencing performance across datasets and sex groups for each model, where the correlations did not change. For example in the case of EEGNet, in the Cho 2017 dataset, the partial correlation yielded a value of *r* = 0.82, confirming that the influence of class distinctiveness on model performance was still significant even after controlling for sex. Similarly, in the Lee 2019 dataset, the partial correlation for all subjects was *r* = 0.33. In all, these results show how spurious correlations between sex and class distinctiveness may lead to erroneous interpretations when analyzing biases (see Table S5).

To delve deeper in the analysis, we next applied mixed-effects models to understand the influence of class distinctiveness, sex, and age on performance metrics, using subjects as groups so that the effects of the training process and model initializations are treated as random effects. When analyzing the CSP+LDA results, we found that class distinctiveness had a positive contribution to performance metrics, as expected. For instance, in the Cho 2017 dataset, the coefficient for class distinctiveness was 0.07 ± 0.01, indicating a significant relationship between class distinctiveness and accuracy. Similarly, sex contributed positively, with a coefficient of 0.09 ± 0.04. These results suggest that both variables significantly influence model performance. Additionally, age had a notable effect, with a coefficient of 0.03 ± 0.002, reinforcing its role in shaping the accuracies of models (see Figure S3 for age distribution in each dataset). Regarding the Lee 2019 dataset, neither sex nor class distinctiveness showed significant effects. In session 1, the coefficient for class distinctiveness was 0.02± 0.01 and for sex it was −0.02 ± 0.03, and in session 2 the coefficients were 0.01 ± 0.01 and −0.02 ± 0.03, respectively. However, age remained significant in both sessions, with a coefficient of 0.03 ± 0.002 in session 1 and of 0.02 ± 0.001 in session 2.

The EEGNet results revealed a more weak interaction between the variables. For the Cho 2017 dataset, class distinctiveness had a coefficient of −0.06± 0.01, while sex did not achieve significance with a coefficient of 0.05± 0.04. As observed in CSP+LDA, age was a strong predictor, with a coefficient of 0.03 ±0.002. For the Lee 2019 session 1 dataset, neither class distinctiveness (0.01± 0.02) nor sex (−0.06 ±0.04) demonstrated statistical significance. Similarly, in Lee 2019 session 2, class distinctiveness had a coefficient of 0.01± 0.02, and sex had −0.06±0.04, with both variables remaining non-significant. Nevertheless, age retained its consistent role as a significant predictor, with coefficients of 0.03 for both sessions. All this information and the analysis for AUC metrics are available in Table S6 and Table S7.

The results highlight the consistent importance of class distinctiveness in CSP+LDA, particularly in Cho 2017, where its effect was statistically significant. However, the effect of class distinctiveness was less evident in EEG-Net, suggesting that DL methods may enhance classification performance even when class distinctiveness is lower. Additionally, the role of sex was less pronounced across both models, with significance observed only in CSP+LDA for Cho 2017.

The comparison underscores the robustness of EEGNet in mitigating the direct influence of demographic variables like sex, while CSP+LDA appears more sensitive to these factors. These findings suggest that while, as expected, class distinctiveness serves as a critical driver of performance, sex itself, when isolated from other confounders, exerts a relatively minor influence in a DL model like EEGNet. A similar tendency was observed for all tested deep learning models, showing that the results obtained are not a consequence of a particular DL architecture but stem from the data itself. Detailed results can be found in Tables Section S2 of the Supplementary material.

## 4 Discussion

While deep learning models (DL) have the potential to improve the efficacy of MI-BCIs, the complexity of such models, which strongly relies on a hierarchy of representations of the data, may lead to biases with respect to demographic attributes. This work focuses on addressing a critical gap regarding the fairness of DL models for MI-BCI, namely the potential *sources* of these biases. In this work we have systematically studied whether DL models generate or amplify biases with respect to the biological sex of the participants in MI-BCI tasks, even when employing a balanced dataset during model training. To that end, we tested model performance in across-subject MI-BCI systems employing both a traditional approach based on statistical machine learning (CSP+LDA), and a variety of Deep Learning models. Although the analysis presented in the main text corresponds to the popular EEGNet architecture, the insights gained were validated on multiple other architectures (presented in the Supplementary Material). Overall, the conclusions derived in this study stem from an analysis of a total of 7 architectures, 38,400 individual models, and 106 subjects from two separate databases.

The results of this study initially showed DL models help the performance of the entire population in an MI-BCI task with respect to the traditional CSP+LDA method, with a consistent (though non statistically significant) trend showing higher female performance. Aware of potential confounders, we resorted to a statistical mixed-effects model to explore how different variables such as sex, class distinctiveness and age contribute to performance metrics. As expected, class distinctiveness appeared as a strong predictor with statistical significance, while sex contributed in a lower proportion, especially in the case of the DL models. This analysis was important for understanding the relationship between the information in the original data and the contributions of different models, aiming to identify whether potential biases are introduced by the DL model or are intrinsic to the data. Notably, while subjects with higher class distinctiveness generally performed better, some participants with lower values still achieved good performance. While class distinctiveness is a valuable metric for approximating the discriminability of patterns related to MI-tasks, it does not fully explain the variability in performance. It is worth mentioning that the employed class distinctiveness metric does not measure the best possible separability of the signals, but represents a lower bound estimate. When this value is high, DL models might not be the best option because improvements are lower at the expense of more computational requirements. Conversely, when class distinctiveness is low, information related to MI tasks may be obscured by other factors, such as protected attributes present in the signal. If protected attributes are present in the data, complex models might inadvertently exploit them, potentially contributing to performance disparities. This may happen due to spurious correlations in the training set that do not generalize to the test set, and may be a particular cause for concern in the case of minorities, where models may sacrifice performance in a small group to increase the overall performance. Furthermore, class distinctiveness may also serve as an indirect indicator of task difficulty, shedding light on why certain models perform differently across subgroups. Our analyses indicate that the gap on performance metrics across females and males might not be due to sex differences themselves. Instead, participants who generate more stable and distinct EEG signals tend to perform better. This conclusion is further supported by a frequency-band ablation study, which demonstrated that model performance remains stable even after removing individual frequency bands (Section S4.3, Table S11).

Therefore, the apparent advantage observed in females in these datasets seems to reflect the presence of subjects with higher MI modulation capabilities, who were predominantly women, rather than biases introduced by the model. This interpretation is strongly supported by additional robustness checks. Specifically, when the EEGNet model was trained directly on the MI data to classify subject sex (Section S4.1 Table S9), performance remained near chance level, confirming the absence of readily exploitable, explicit sex features. While this result would seem to deviate from recent successful sex classification reports, previous work performed classification using datasets that involve passive states (e.g., watching movies [14] or resting state with closed/opened eyes [15, 16, 17, 18]). These passive tasks could reveal stable physiological differences, whereas the active cognitive MI-BCI task may mask these sex-related neural features. Also, passive tasks allow for longer acquisition sessions per subject, providing a large volume of data making the training process of DL models more efficient. Collecting such extensive data is often impractical in MI-BCI tasks due to the constraints of user fatigue and the demand for sustained high cognitive effort. Furthermore, exploratory experiments with severe sex imbalances (ranging from 25% to 75% female representation) demonstrated that the core performance disparity remained stable (Section S4.2 Table S10). Why females show better performance in terms of class distinctiveness with these datasets, and whether this trend generalizes to other datasets remain open questions to be investigated.

To address these considerations comprehensively, there is a pressing need for more diverse BCI datasets specifying metadata information, but also for the development of innovative experimental procedures. For example, prior studies have demonstrated that the interaction between the sex of the participant and the sex of the experimenter may also play a significant role [34]. So, future datasets should incorporate detailed metadata from both experimenters and participants, including socioeconomic status, ethnicity, geographic region, and physical traits (e.g., hair type). Moreover, the level of expertise or previous exposure to BCI tasks could be essential to understand inter-subject variability. This broader range of metadata would allow for a more exhaustive and intersectional analysis of how individual differences influence EEG patterns and model performance.

We have also highlighted how the LOSO approach, which is commonly employed as a validation scheme, compromises the assumptions of statistical independence, limiting the possibility to apply traditional hypothesis tests to establish statistical significance. In this work we overcome this limitation using partial correlations and mixed-effect models, but one alternative solution for larger databases would be to train models based on one subset of the database and then test on multiple individual users not included in that subset of the database, potentially allowing for statistical tests to assess performance differences. For a fixed dataset size, this would significantly reduce statistical power and model performance. This approach would hence require counting with larger databases, or testing across datasets (which is not without challenges). Future work should include these types of analyses. In addition, promising avenues of research include the systematic exploration of multiple classification algorithms, including classical ML methods, together with this and other DL models. Furthermore, longitudinal studies could investigate how repeated BCI training influences performance and class distinctiveness over time.

To conclude, our findings suggest that the predictive DL model is not in itself responsible for sex-related biases in the absence of data imbalance in terms of sex in the training population, which is a significant consideration for real-world BCI applications. Additionally, while participants with higher MI capability seem not to require complex models, those with lower MI-BCI skills substantially benefit from more sophisticated models such DL architectures. These combined observations are highly encouraging for the use of DL methods for MI-BCI. Indeed, ensuring that decoding algorithms do not introduce biases (or that they can eventually be intervened to mitigate them), is a key step towards improved access to fair BCI systems. We believe that an analysis like the one performed here, which goes beyond standard evaluation of performance, or even classical stratification should become standard practices to reach this goal. This is especially critical in rehabilitation contexts, where the effectiveness of a system should not disproportionately favor certain groups over others. Enabling equitable performance across diverse populations would enhance the accessibility and reliability of BCI technologies for all users.

## 5 Acknowledgments

This work was supported by Argentina’s National Scientific and Technical Research Council (CONICET), who covered all researchers’ salaries and scholarships.

