## Supplementary Material for "Exploring sex-related Biases in Deep Learning Models for Motor Imagery Brain-Computer Interfaces"

#### S1 Distribution of class distinctiveness

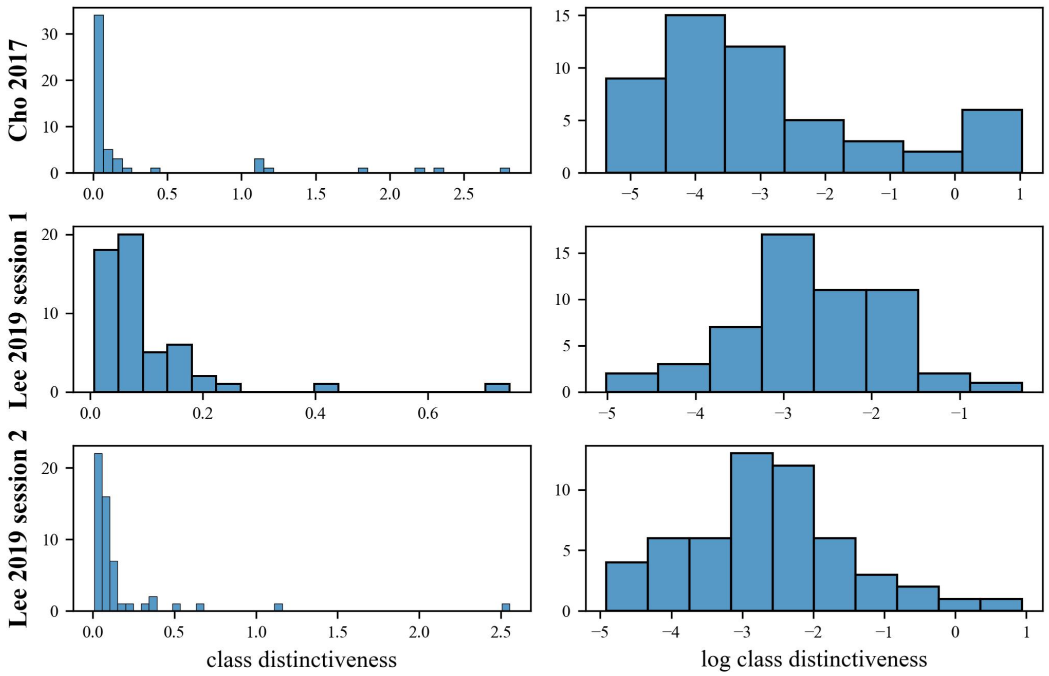


Figure S1: Histogram of class distinctiveness for the different datasets and sessions (row) in linear scale (left column) and logarithmic scale (right column).

#### S2 Validation of EEGNet Results with Multiple Models and Methods

This section presents the results obtained from the training process of various Deep Learning (DL) models. The general training procedure followed the methodology detailed in the main paper, but with specific differences in the number of trained models and random seeds used:

- For EEGNet, we trained 100 models per subject (5 model initialization seeds $\times$ 20 data split seeds).
- For the remaining DL models, we trained 20 models per subject (2 model initialization seeds $\times$ 10 data split seeds).
- For the CSP+LDA method, we trained 20 models per subject (using 20 data split seeds).

The subsequent subsections detail the specific results achieved in each conducted analysis.

##### S2.1 Comparison of Model Performance

Table S1: Performance comparison in terms of accuracy of different models for MI classification, showing the global metric and segregated by sex. A Mann-Whitney U test was applied to assess differences between female and male subjects, and the resulting p-values were adjusted using the Bonferroni correction due to the number of comparisons ($n = 8$).

| **Dataset** | **Model** | **Global Acc.** | **Females Acc.** | **Males Acc.** | **p-value** |
|---|---|---|---|---|---|
| Cho 2017 | EEGNetv4 | $0.71 \pm 0.13$ | $0.75 \pm 0.13$ | $0.68 \pm 0.12$ | 0.44 |
| | Deep4Net | $0.71 \pm 0.13$ | $0.75 \pm 0.13$ | $0.68 \pm 0.13$ | 0.54 |
| | ShallowFBCSPNet | $0.70 \pm 0.13$ | $0.74 \pm 0.13$ | $0.68 \pm 0.13$ | 0.92 |
| | HybridNet | $0.69 \pm 0.13$ | $0.73 \pm 0.13$ | $0.67 \pm 0.13$ | 0.70 |
| | FBCNet | $0.59 \pm 0.08$ | $0.62 \pm 0.11$ | $0.57 \pm 0.06$ | 1.00 |
| | CTNet | $0.70 \pm 0.13$ | $0.74 \pm 0.13$ | $0.68 \pm 0.13$ | 0.71 |
| | ATCNet | $0.70 \pm 0.13$ | $0.75 \pm 0.13$ | $0.68 \pm 0.13$ | 0.38 |
| | CSP+LDA | $0.59 \pm 0.15$ | $0.62 \pm 0.14$ | $0.58 \pm 0.15$ | 0.99 |
| Lee 2019 Session 1 | EEGNetv4 | $0.71 \pm 0.12$ | $0.74 \pm 0.11$ | $0.69 \pm 0.12$ | 1.00 |
| | Deep4Net | $0.71 \pm 0.12$ | $0.74 \pm 0.11$ | $0.69 \pm 0.11$ | 0.86 |
| | ShallowFBCSPNet | $0.72 \pm 0.11$ | $0.75 \pm 0.10$ | $0.69 \pm 0.12$ | 0.53 |
| | HybridNet | $0.71 \pm 0.12$ | $0.74 \pm 0.11$ | $0.68 \pm 0.12$ | 0.44 |
| | FBCNet | $0.62 \pm 0.10$ | $0.64 \pm 0.09$ | $0.60 \pm 0.11$ | 0.25 |
| | CTNet | $0.71 \pm 0.11$ | $0.73 \pm 0.10$ | $0.69 \pm 0.12$ | 1.00 |
| | ATCNet | $0.70 \pm 0.12$ | $0.73 \pm 0.11$ | $0.69 \pm 0.12$ | 1.00 |
| | CSP+LDA | $0.59 \pm 0.08$ | $0.59 \pm 0.09$ | $0.58 \pm 0.07$ | 1.00 |
| Lee 2019 Session 2 | EEGNetv4 | $0.71 \pm 0.12$ | $0.74 \pm 0.11$ | $0.69 \pm 0.12$ | 1.00 |
| | Deep4Net | $0.71 \pm 0.12$ | $0.75 \pm 0.11$ | $0.68 \pm 0.12$ | 0.44 |
| | ShallowFBCSPNet | $0.71 \pm 0.12$ | $0.75 \pm 0.11$ | $0.69 \pm 0.12$ | 0.62 |
| | HybridNet | $0.71 \pm 0.12$ | $0.74 \pm 0.10$ | $0.69 \pm 0.12$ | 1.00 |
| | FBCNet | $0.62 \pm 0.11$ | $0.65 \pm 0.12$ | $0.61 \pm 0.10$ | 0.99 |
| | CTNet | $0.70 \pm 0.11$ | $0.73 \pm 0.10$ | $0.68 \pm 0.12$ | 1.00 |
| | ATCNet | $0.70 \pm 0.11$ | $0.73 \pm 0.10$ | $0.68 \pm 0.12$ | 1.00 |
| | CSP+LDA | $0.55 \pm 0.08$ | $0.56 \pm 0.08$ | $0.54 \pm 0.08$ | 1.00 |

Table S2: Performance comparison of different models for MI classification in terms of AUC, showing the global metric and segregated by sex. A Mann-Whitney U test was applied to assess differences between female and male subjects, and the resulting p-values were adjusted using the Bonferroni correction due to the number of comparisons ($n = 8$).

| **Dataset** | **Model** | **Global AUC** | **Females AUC** | **Males AUC** | **p-value** |
|---|---|---|---|---|---|
| Cho 2017 | EEGNetv4 | $0.77 \pm 0.14$ | $0.82 \pm 0.13$ | $0.74 \pm 0.13$ | 0.34 |
| | Deep4Net | $0.77 \pm 0.14$ | $0.82 \pm 0.13$ | $0.74 \pm 0.13$ | 0.34 |
| | ShallowFBCSPNet | $0.77 \pm 0.14$ | $0.82 \pm 0.14$ | $0.74 \pm 0.13$ | 0.67 |
| | HybridNet | $0.76 \pm 0.14$ | $0.81 \pm 0.14$ | $0.73 \pm 0.13$ | 0.46 |
| | FBCNet | $0.63 \pm 0.11$ | $0.67 \pm 0.13$ | $0.60 \pm 0.08$ | 0.92 |
| | CTNet | $0.76 \pm 0.14$ | $0.82 \pm 0.14$ | $0.73 \pm 0.13$ | 0.42 |
| | ATCNet | $0.76 \pm 0.14$ | $0.82 \pm 0.13$ | $0.73 \pm 0.13$ | 0.25 |
| | CSP+LDA | $0.62 \pm 0.18$ | $0.66 \pm 0.18$ | $0.60 \pm 0.17$ | 1.00 |
| Lee 2019 Session 1 | EEGNetv4 | $0.78 \pm 0.13$ | $0.82 \pm 0.11$ | $0.75 \pm 0.14$ | 0.49 |
| | Deep4Net | $0.79 \pm 0.13$ | $0.83 \pm 0.11$ | $0.75 \pm 0.14$ | 0.37 |
| | ShallowFBCSPNet | $0.79 \pm 0.13$ | $0.84 \pm 0.10$ | $0.76 \pm 0.14$ | 0.25 |
| | HybridNet | $0.78 \pm 0.14$ | $0.83 \pm 0.11$ | $0.75 \pm 0.14$ | 0.21 |
| | FBCNet | $0.68 \pm 0.13$ | $0.72 \pm 0.12$ | $0.65 \pm 0.14$ | 0.09 |
| | CTNet | $0.78 \pm 0.13$ | $0.82 \pm 0.11$ | $0.75 \pm 0.14$ | 0.69 |
| | ATCNet | $0.77 \pm 0.13$ | $0.81 \pm 0.11$ | $0.74 \pm 0.14$ | 0.69 |
| | CSP+LDA | $0.64 \pm 0.11$ | $0.65 \pm 0.12$ | $0.62 \pm 0.10$ | 1.00 |
| Lee 2019 Session 2 | EEGNetv4 | $0.78 \pm 0.13$ | $0.82 \pm 0.11$ | $0.75 \pm 0.14$ | 0.80 |
| | Deep4Net | $0.78 \pm 0.13$ | $0.83 \pm 0.10$ | $0.75 \pm 0.14$ | 0.18 |
| | ShallowFBCSPNet | $0.79 \pm 0.13$ | $0.83 \pm 0.10$ | $0.75 \pm 0.14$ | 0.27 |
| | HybridNet | $0.78 \pm 0.13$ | $0.82 \pm 0.10$ | $0.75 \pm 0.14$ | 0.57 |
| | FBCNet | $0.68 \pm 0.14$ | $0.72 \pm 0.14$ | $0.66 \pm 0.14$ | 0.55 |
| | CTNet | $0.77 \pm 0.13$ | $0.82 \pm 0.10$ | $0.74 \pm 0.14$ | 0.86 |
| | ATCNet | $0.77 \pm 0.13$ | $0.82 \pm 0.11$ | $0.74 \pm 0.14$ | 0.74 |
| | CSP+LDA | $0.58 \pm 0.13$ | $0.58 \pm 0.13$ | $0.57 \pm 0.13$ | 1.00 |

#### S2.2 Performance and Class Distinctiveness correlations for all tested methods

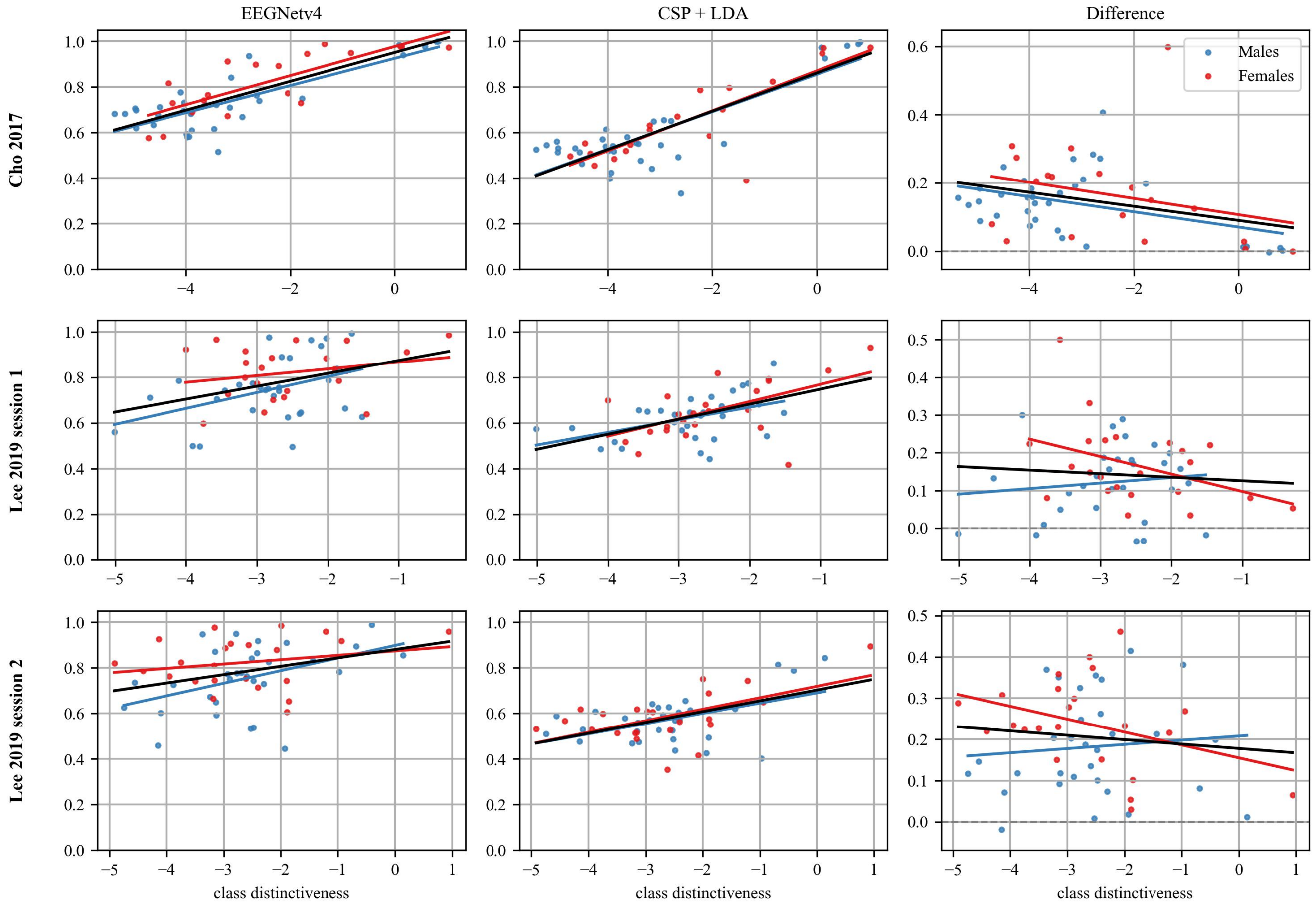


Figure S2: Scatterplots of AUC vs. class distinctiveness for both databases. Each row corresponds to a specific database, and the columns represent the AUC performance of the EEGNet model, CSP+LDA, and their differences. Each dot represents the average performance across all models trained to test for a single subject. Data points for males are shown in blue, while those for females are in red. Straight lines indicate regression trends: black lines represent regression across all data (not segregated by sex), and colored lines correspond to sex-specific regressions.

Table S3: Correlation metrics between accuracy and class distinctiveness for all datasets. The correlation coefficient ($r$) and mean squared error (MSE) for all models are presented, including overall results and sex-segregated performance.

| **Model** | **Data** | **Cho 2017** | | **Lee 2019 session 1** | | **Lee 2019 session 2** | |
|---|---|---|---|---|---|---|---|
| | | $r$ | **MSE** | $r$ | **MSE** | $r$ | **MSE** |
| EEGNet | all data | 0.83 | $5.16 \times 10^{-3}$ | 0.36 | $1.14 \times 10^{-2}$ | 0.32 | $1.16 \times 10^{-2}$ |
| | F | 0.79 | $6.20 \times 10^{-3}$ | 0.22 | $1.07 \times 10^{-2}$ | 0.24 | $1.01 \times 10^{-2}$ |
| | M | 0.85 | $4.25 \times 10^{-3}$ | 0.42 | $1.10 \times 10^{-2}$ | 0.40 | $1.14 \times 10^{-2}$ |
| Deep4Net | all data | 0.83 | $5.16 \times 10^{-3}$ | 0.40 | $1.04 \times 10^{-2}$ | 0.29 | $1.21 \times 10^{-2}$ |
| | F | 0.81 | $5.59 \times 10^{-3}$ | 0.31 | $9.92 \times 10^{-3}$ | 0.33 | $9.09 \times 10^{-3}$ |
| | M | 0.83 | $4.69 \times 10^{-3}$ | 0.43 | $1.01 \times 10^{-2}$ | 0.29 | $1.25 \times 10^{-2}$ |
| FBCNet | all data | 0.32 | $5.50 \times 10^{-3}$ | 0.25 | $8.80 \times 10^{-3}$ | 0.22 | $1.05 \times 10^{-2}$ |
| | F | 0.66 | $5.84 \times 10^{-3}$ | 0.31 | $6.65 \times 10^{-3}$ | 0.37 | $1.05 \times 10^{-2}$ |
| | M | -0.10 | $2.75 \times 10^{-3}$ | 0.16 | $9.69 \times 10^{-3}$ | 0.10 | $9.54 \times 10^{-3}$ |
| CTNet | all data | 0.84 | $5.10 \times 10^{-3}$ | 0.39 | $1.03 \times 10^{-2}$ | 0.32 | $1.07 \times 10^{-2}$ |
| | F | 0.81 | $5.74 \times 10^{-3}$ | 0.31 | $8.89 \times 10^{-3}$ | 0.33 | $8.15 \times 10^{-3}$ |
| | M | 0.84 | $4.56 \times 10^{-3}$ | 0.41 | $1.08 \times 10^{-2}$ | 0.34 | $1.12 \times 10^{-2}$ |
| ATCNet | all data | 0.84 | $4.84 \times 10^{-3}$ | 0.38 | $1.04 \times 10^{-2}$ | 0.33 | $1.04 \times 10^{-2}$ |
| | F | 0.82 | $5.24 \times 10^{-3}$ | 0.24 | $9.32 \times 10^{-3}$ | 0.30 | $8.21 \times 10^{-3}$ |
| | M | 0.85 | $4.40 \times 10^{-3}$ | 0.44 | $1.05 \times 10^{-2}$ | 0.40 | $1.08 \times 10^{-2}$ |
| HybridNet | all data | 0.84 | $5.06 \times 10^{-3}$ | 0.39 | $1.07 \times 10^{-2}$ | 0.33 | $1.09 \times 10^{-2}$ |
| | F | 0.84 | $4.65 \times 10^{-3}$ | 0.28 | $9.70 \times 10^{-3}$ | 0.38 | $8.31 \times 10^{-3}$ |
| | M | 0.82 | $5.16 \times 10^{-3}$ | 0.42 | $1.03 \times 10^{-2}$ | 0.33 | $1.15 \times 10^{-2}$ |
| ShallowFBCSPNet | all data | 0.83 | $5.45 \times 10^{-3}$ | 0.41 | $1.03 \times 10^{-2}$ | 0.27 | $1.19 \times 10^{-2}$ |
| | F | 0.81 | $5.99 \times 10^{-3}$ | 0.35 | $8.16 \times 10^{-3}$ | 0.36 | $9.08 \times 10^{-3}$ |
| | M | 0.83 | $4.98 \times 10^{-3}$ | 0.41 | $1.10 \times 10^{-2}$ | 0.25 | $1.23 \times 10^{-2}$ |
| CSP+LDA | all data | 0.86 | $5.45 \times 10^{-3}$ | 0.52 | $4.16 \times 10^{-3}$ | 0.55 | $2.77 \times 10^{-3}$ |
| | F | 0.85 | $5.28 \times 10^{-3}$ | 0.60 | $4.57 \times 10^{-3}$ | 0.59 | $2.63 \times 10^{-3}$ |
| | M | 0.87 | $5.49 \times 10^{-3}$ | 0.42 | $3.70 \times 10^{-3}$ | 0.53 | $2.76 \times 10^{-3}$ |

Table S4: Correlation metrics between AUC and class distinctiveness for all datasets. The correlation coefficient ($r$) and mean squared error (MSE) for all models are presented, including overall results and sex-segregated performance.

| **Model** | **Data** | **Cho 2017** | | **Lee 2019 session 1** | | **Lee 2019 session 2** | |
|---|---|---|---|---|---|---|---|
| | | $r$ | **MSE** | $r$ | **MSE** | $r$ | **MSE** |
| EEGNet | all data | 0.81 | $6.40 \times 10^{-3}$ | 0.37 | $1.53 \times 10^{-2}$ | 0.32 | $1.58 \times 10^{-2}$ |
| | F | 0.79 | $6.53 \times 10^{-3}$ | 0.24 | $1.17 \times 10^{-2}$ | 0.22 | $1.11 \times 10^{-2}$ |
| | M | 0.81 | $5.73 \times 10^{-3}$ | 0.41 | $1.59 \times 10^{-2}$ | 0.43 | $1.64 \times 10^{-2}$ |
| Deep4Net | all data | 0.80 | $6.53 \times 10^{-3}$ | 0.39 | $1.45 \times 10^{-2}$ | 0.27 | $1.62 \times 10^{-2}$ |
| | F | 0.81 | $6.23 \times 10^{-3}$ | 0.32 | $1.03 \times 10^{-2}$ | 0.23 | $9.84 \times 10^{-3}$ |
| | M | 0.79 | $6.09 \times 10^{-3}$ | 0.39 | $1.59 \times 10^{-2}$ | 0.35 | $1.71 \times 10^{-2}$ |
| FBCNet | all data | 0.28 | $1.04 \times 10^{-2}$ | 0.29 | $1.56 \times 10^{-2}$ | 0.21 | $1.86 \times 10^{-2}$ |
| | F | 0.62 | $1.02 \times 10^{-2}$ | 0.43 | $1.00 \times 10^{-2}$ | 0.38 | $1.66 \times 10^{-2}$ |
| | M | -0.09 | $6.39 \times 10^{-3}$ | 0.15 | $1.78 \times 10^{-2}$ | 0.09 | $1.77 \times 10^{-2}$ |
| CTNet | all data | 0.81 | $6.51 \times 10^{-3}$ | 0.39 | $1.46 \times 10^{-2}$ | 0.30 | $1.57 \times 10^{-2}$ |
| | F | 0.81 | $6.09 \times 10^{-3}$ | 0.33 | $1.04 \times 10^{-2}$ | 0.24 | $9.60 \times 10^{-3}$ |
| | M | 0.80 | $6.22 \times 10^{-3}$ | 0.40 | $1.63 \times 10^{-2}$ | 0.38 | $1.72 \times 10^{-2}$ |
| ATCNet | all data | 0.83 | $6.07 \times 10^{-3}$ | 0.38 | $1.46 \times 10^{-2}$ | 0.32 | $1.54 \times 10^{-2}$ |
| | F | 0.83 | $5.53 \times 10^{-3}$ | 0.25 | $1.11 \times 10^{-2}$ | 0.23 | $9.16 \times 10^{-3}$ |
| | M | 0.82 | $5.77 \times 10^{-3}$ | 0.42 | $1.56 \times 10^{-2}$ | 0.43 | $1.67 \times 10^{-2}$ |
| HybridNet | all data | 0.81 | $6.70 \times 10^{-3}$ | 0.39 | $1.51 \times 10^{-2}$ | 0.31 | $1.52 \times 10^{-2}$ |
| | F | 0.83 | $5.69 \times 10^{-3}$ | 0.31 | $1.00 \times 10^{-2}$ | 0.27 | $9.46 \times 10^{-3}$ |
| | M | 0.78 | $6.74 \times 10^{-3}$ | 0.40 | $1.65 \times 10^{-2}$ | 0.39 | $1.66 \times 10^{-2}$ |
| ShallowFBCSPNet | all data | 0.80 | $6.92 \times 10^{-3}$ | 0.40 | $1.46 \times 10^{-2}$ | 0.26 | $1.58 \times 10^{-2}$ |
| | F | 0.81 | $6.67 \times 10^{-3}$ | 0.37 | $8.37 \times 10^{-3}$ | 0.24 | $9.15 \times 10^{-3}$ |
| | M | 0.78 | $6.52 \times 10^{-3}$ | 0.38 | $1.71 \times 10^{-2}$ | 0.32 | $1.77 \times 10^{-2}$ |
| CSP+LDA | all data | 0.83 | $9.66 \times 10^{-3}$ | 0.53 | $8.49 \times 10^{-3}$ | 0.52 | $8.28 \times 10^{-3}$ |
| | F | 0.82 | $9.90 \times 10^{-3}$ | 0.56 | $1.02 \times 10^{-2}$ | 0.57 | $8.41 \times 10^{-3}$ |
| | M | 0.83 | $9.50 \times 10^{-3}$ | 0.47 | $7.07 \times 10^{-3}$ | 0.49 | $8.07 \times 10^{-3}$ |

##### S2.3 Partial Correlation Tables

Table S5: Partial correlation coefficients ($r$) between performance metrics (Accuracy and AUC) and the logarithm of class distinctiveness, using sex as a covariate. The table shows the correlation coefficient for each dataset and model.

| **Metric** | **Model** | **Cho 2017** | **Lee 2019 session 1** | **Lee 2019 session 2** |
|---|---|---|---|---|
| | | **r** | **r** | **r** |
| **Accuracy** | EEGNet | 0.82 | 0.33 | 0.33 |
| | Deep4Net | 0.82 | 0.37 | 0.31 |
| | FBCNet | 0.27 | 0.22 | 0.23 |
| | CTNet | 0.83 | 0.37 | 0.33 |
| | ATCNet | 0.84 | 0.36 | 0.35 |
| | HybridNet | 0.83 | 0.36 | 0.35 |
| | ShallowFBCSPNet | 0.82 | 0.38 | 0.29 |
| | CSP+LDA | 0.86 | 0.51 | 0.56 |
| **AUC** | EEGNet | 0.80 | 0.34 | 0.34 |
| | Deep4Net | 0.80 | 0.36 | 0.30 |
| | FBCNet | 0.23 | 0.26 | 0.22 |
| | CTNet | 0.80 | 0.36 | 0.32 |
| | ATCNet | 0.82 | 0.35 | 0.35 |
| | HybridNet | 0.80 | 0.36 | 0.34 |
| | ShallowFBCSPNet | 0.79 | 0.37 | 0.28 |
| | CSP+LDA | 0.83 | 0.52 | 0.52 |

#### S2.4 Mixed-Effects Models

Table S6: Mixed-Effects Models for the influence on accuracy of class distinctiveness, sex, and age, using subjects as groups so that the effects of the training process and model initializations are treated as random effects.

| Variable | Model | Cho 2017 Coefficient | Lee 2019 session 1 Coefficient | Lee 2019 session 2 Coefficient |
|---|---|---|---|---|
| class distinctiveness | EEGNet | $0.06 \pm 0.01$ | $0.01 \pm 0.02$ | $0.01 \pm 0.02$ |
| | Deep4Net | $0.06 \pm 0.01$ | $0.02 \pm 0.02$ | $0.01 \pm 0.02$ |
| | ShallowFBCSPNet | $0.06 \pm 0.01$ | $0.02 \pm 0.02$ | $0.01 \pm 0.02$ |
| | HybridNet | $0.06 \pm 0.01$ | $0.01 \pm 0.02$ | $0.01 \pm 0.02$ |
| | FBCNet | $0.01 \pm 0.01$ | $-0.01 \pm 0.02$ | $0.00 \pm 0.01$ |
| | CTNet | $0.06 \pm 0.01$ | $0.02 \pm 0.02$ | $0.01 \pm 0.02$ |
| | ATCNet | $0.06 \pm 0.01$ | $0.02 \pm 0.02$ | $0.01 \pm 0.02$ |
| | CSP+LDA | $0.07 \pm 0.01$ | $0.02 \pm 0.02$ | $0.01 \pm 0.01$ |
| sex | EEGNet | $0.05 \pm 0.04$ | $-0.06 \pm 0.04$ | $-0.06 \pm 0.04$ |
| | Deep4Net | $0.06 \pm 0.04$ | $-0.06 \pm 0.04$ | $-0.08 \pm 0.04$ |
| | ShallowFBCSPNet | $0.06 \pm 0.04$ | $-0.07 \pm 0.04$ | $-0.07 \pm 0.04$ |
| | HybridNet | $0.06 \pm 0.04$ | $-0.07 \pm 0.04$ | $-0.07 \pm 0.04$ |
| | FBCNet | $0.02 \pm 0.03$ | $-0.06 \pm 0.04$ | $-0.05 \pm 0.04$ |
| | CTNet | $0.06 \pm 0.04$ | $-0.06 \pm 0.04$ | $-0.07 \pm 0.04$ |
| | ATCNet | $0.06 \pm 0.04$ | $-0.05 \pm 0.04$ | $-0.06 \pm 0.04$ |
| | CSP+LDA | $0.09 \pm 0.04$ | $-0.02 \pm 0.03$ | $-0.02 \pm 0.03$ |
| age | EEGNet | $0.03 \pm 0.00$ | $0.03 \pm 0.00$ | $0.03 \pm 0.00$ |
| | Deep4Net | $0.03 \pm 0.00$ | $0.03 \pm 0.00$ | $0.03 \pm 0.00$ |
| | ShallowFBCSPNet | $0.03 \pm 0.00$ | $0.03 \pm 0.00$ | $0.03 \pm 0.00$ |
| | HybridNet | $0.03 \pm 0.00$ | $0.03 \pm 0.00$ | $0.03 \pm 0.00$ |
| | FBCNet | $0.02 \pm 0.00$ | $0.03 \pm 0.00$ | $0.03 \pm 0.00$ |
| | CTNet | $0.03 \pm 0.00$ | $0.03 \pm 0.00$ | $0.03 \pm 0.00$ |
| | ATCNet | $0.03 \pm 0.00$ | $0.03 \pm 0.00$ | $0.03 \pm 0.00$ |
| | CSP+LDA | $0.03 \pm 0.00$ | $0.03 \pm 0.00$ | $0.02 \pm 0.00$ |

Table S7: Mixed-Effect Models for the influence on AUC of class distinctiveness, sex, and age, using subjects as groups so that the effects of the training process and model initializations are treated as random effects.

| **Variable** | **Model** | **Cho 2017 Coefficient** | **Lee 2019 session 1 Coefficient** | **Lee 2019 session 2 Coefficient** |
|---|---|---|---|---|
| class distinctiveness | EEGNet | $0.06 \pm 0.01$ | $0.02 \pm 0.02$ | $0.01 \pm 0.02$ |
| | Deep4Net | $0.06 \pm 0.01$ | $0.02 \pm 0.02$ | $0.01 \pm 0.02$ |
| | ShallowFBCSPNet | $0.06 \pm 0.01$ | $0.02 \pm 0.02$ | $0.01 \pm 0.02$ |
| | HybridNet | $0.06 \pm 0.01$ | $0.02 \pm 0.02$ | $0.01 \pm 0.02$ |
| | FBCNet | $0.01 \pm 0.01$ | $0.01 \pm 0.02$ | $0.01 \pm 0.02$ |
| | CTNet | $0.06 \pm 0.01$ | $0.02 \pm 0.02$ | $0.01 \pm 0.02$ |
| | ATCNet | $0.06 \pm 0.01$ | $0.02 \pm 0.02$ | $0.01 \pm 0.02$ |
| | CSP+LDA | $0.08 \pm 0.01$ | $0.04 \pm 0.02$ | $0.03 \pm 0.01$ |
| sex | EEGNet | $0.05 \pm 0.05$ | $-0.09 \pm 0.04$ | $-0.09 \pm 0.04$ |
| | Deep4Net | $0.05 \pm 0.05$ | $-0.09 \pm 0.04$ | $-0.10 \pm 0.04$ |
| | ShallowFBCSPNet | $0.05 \pm 0.05$ | $-0.10 \pm 0.04$ | $-0.10 \pm 0.04$ |
| | HybridNet | $0.05 \pm 0.05$ | $-0.10 \pm 0.04$ | $-0.09 \pm 0.04$ |
| | FBCNet | $0.01 \pm 0.04$ | $-0.09 \pm 0.04$ | $-0.08 \pm 0.05$ |
| | CTNet | $0.05 \pm 0.05$ | $-0.09 \pm 0.04$ | $-0.09 \pm 0.05$ |
| | ATCNet | $0.05 \pm 0.05$ | $-0.08 \pm 0.04$ | $-0.09 \pm 0.04$ |
| | CSP+LDA | $0.08 \pm 0.05$ | $-0.04 \pm 0.04$ | $-0.02 \pm 0.04$ |
| age | EEGNet | $0.04 \pm 0.00$ | $0.04 \pm 0.00$ | $0.04 \pm 0.00$ |
| | Deep4Net | $0.04 \pm 0.00$ | $0.04 \pm 0.00$ | $0.04 \pm 0.00$ |
| | ShallowFBCSPNet | $0.04 \pm 0.00$ | $0.04 \pm 0.00$ | $0.04 \pm 0.00$ |
| | HybridNet | $0.04 \pm 0.00$ | $0.04 \pm 0.00$ | $0.04 \pm 0.00$ |
| | FBCNet | $0.03 \pm 0.00$ | $0.03 \pm 0.00$ | $0.03 \pm 0.00$ |
| | CTNet | $0.04 \pm 0.00$ | $0.04 \pm 0.00$ | $0.04 \pm 0.00$ |
| | ATCNet | $0.04 \pm 0.00$ | $0.04 \pm 0.00$ | $0.04 \pm 0.00$ |
| | CSP+LDA | $0.03 \pm 0.00$ | $0.03 \pm 0.00$ | $0.03 \pm 0.00$ |

#### S3 Databases Characterization

This section provides an analysis characterizing the demographic composition of each dataset. Specifically, we examined the age distribution and the proportion of the BCI-naïve population. These data are presented in Figure S3 (age distribution) and Table S8 (BCI experience distribution).

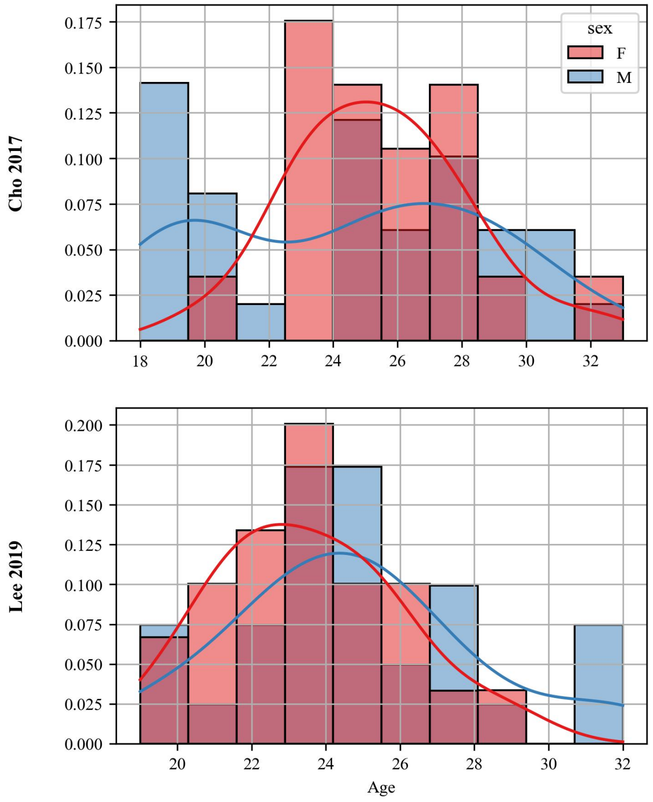


Figure S3: Age distribution segregated by sex. Each row corresponds to a specific database.

Table S8: Participants' Prior BCI Experience. Details on the number of subjects categorized as BCI-naïve (0 prior sessions) or experienced ($\geq$ 1 prior session), segregated by dataset and biological sex. This data confirms the study was conducted on a predominantly BCI-naïve population.

| **Dataset** | **Sex** | **Total Subjects** | **Naïve Subjects** | **Experienced Subjects** | **Prior Session Range** |
|---|---|---|---|---|---|
| Cho 2017 | M | 33 | 28 | 5 | 1 to 10 |
| | F | 19 | 19 | 0 | - |
| Lee 2019 | M | 31 | 20 | 11 | 1 to 4 |
| | F | 23 | 14 | 9 | 1 to 5 |

### S4 Further Analysis into the Sex-related Sources of Bias

#### S4.1 Sex classification

To investigate whether sex-specific features can be extracted by these models and whether features learned for Motor Imagery (MI) Brain-Computer Interface (BCI) classification also encode sex information, we evaluated two distinct scenarios:

- Sex Classification Training from Scratch (directly from the EEG signal): We trained the entire EEGNet architecture from random initialization to classify sex using the MI-EEG input data. This can potentially allow the model to learn sex-discriminative representations directly from the data.

- Transfer Learning for Sex Classification: For each model previously trained for MI-BCI classification, we froze all layers except the final classification layer. We then re-trained only this classification layer for the task of sex prediction. This approach assesses whether the feature representations learned by the model for MI-BCI prediction contain enough information to distinguish between sexes.

In both cases, we followed again a LOSO scheme balanced by sex in both training and validation sets. Results show that sex was not readily classifiable by our models beyond chance level (see Table S9), each result was not significant according to a wilcoxon single rank hypothesis test to compare if the performance of each population was higher than the chance level of 0.5.

Table S9: Sex classification accuracy performance using EEGNet on MI-EEG data. Results show the mean ± standard deviation across subjects from the Leave-One-Subject-Out (LOSO) scheme.

| **Dataset** | **Training from Scratch** | **Transfer Learning** |
|---|---|---|
| Cho2017 | $0.55 \pm 0.33$ | $0.59 \pm 0.30$ |
| Lee2019 session 1 | $0.51 \pm 0.30$ | $0.55 \pm 0.28$ |
| Lee2019 session 2 | $0.50 \pm 0.28$ | $0.55 \pm 0.29$ |

#### S4.2 Robustness to Data Imbalance in Terms of Sex

To assess the model's robustness and understand how particular data distributions might amplify or mask performance disparities, we conducted an exploratory analysis using imbalanced training datasets in terms of sex. We studied three imbalanced training scenarios by varying the ratio of female subjects within the training and validation sets: 25%, 50% (balanced as a control), and 75% female representation. The resulting global and group-specific accuracies for these three imbalance ratios are presented in Table S10. For the most part we see no significant differences in the performance of the models between the two sub-populations.

Table S10: EEGNet accuracy across three different female representation ratios in the training and validation datasets. A Mann-Whitney U test was applied to assess differences between female and male subjects.

| **Dataset** | **Female Ratio** | **Global Acc.** | **Females Acc.** | **Males Acc.** | **Difference (F-M)** |
|---|---|---|---|---|---|
| Cho 2017 | 25% | $0.69 \pm 0.13$ | $0.74 \pm 0.13$ | $0.67 \pm 0.13$ | $0.07^*$ |
| | 50% | $0.70 \pm 0.13$ | $0.74 \pm 0.13$ | $0.67 \pm 0.13$ | 0.07 |
| | 75% | $0.70 \pm 0.13$ | $0.75 \pm 0.13$ | $0.68 \pm 0.12$ | 0.07 |
| Lee 2019 session 1 | 25% | $0.71 \pm 0.12$ | $0.73 \pm 0.11$ | $0.69 \pm 0.12$ | 0.05 |
| | 50% | $0.70 \pm 0.12$ | $0.72 \pm 0.11$ | $0.68 \pm 0.12$ | 0.04 |
| | 75% | $0.70 \pm 0.12$ | $0.73 \pm 0.10$ | $0.68 \pm 0.12$ | 0.05 |
| Lee 2019 session 2 | 25% | $0.70 \pm 0.11$ | $0.72 \pm 0.11$ | $0.68 \pm 0.11$ | 0.04 |
| | 50% | $0.70 \pm 0.12$ | $0.72 \pm 0.11$ | $0.69 \pm 0.12$ | 0.03 |
| | 75% | $0.70 \pm 0.12$ | $0.73 \pm 0.11$ | $0.68 \pm 0.12$ | 0.05 |

##### S4.3 Ablation Study in terms of Frequency-Bands

To asses if model performance relies heavily on certain frequency bands we performed frequency ablation studies. We tested the trained models with testing data filtered using a bandstop filter in different frequency bands: Delta ($< 4$ Hz), Theta ($4 - 8$ Hz), Alpha ($8 - 13$ Hz), Beta ($13 - 30$ Hz), and Gamma ($> 30$ Hz). The results, presented in Table S11, show that the elimination of individual frequency bands did not substantially alter the performance of the models.

Table S11: Impact of Frequency Band Ablation on model performance for MI classification. A Wilcoxon test was done to asses difference between the baseline result vs the ablated results, and the resulting p-values were adjusted using the Bonferroni correction due to the number of comparisons ($n = 5$).

| **Dataset** | **Global Acc.** | **Frequency-Band ablated** | **Global Acc.** | **Difference** | **p-value** |
|---|---|---|---|---|---|
| Cho 2017 | 0.71 ± 0.13 | Alpha | 0.69 ± 0.14 | 0.01 | 0.02 |
| | | Beta | 0.70 ± 0.13 | 0.01 | 1.00 |
| | | Delta | 0.70 ± 0.13 | 0.01 | 0.04 |
| | | Gamma | 0.71 ± 0.13 | 0.00 | 1.00 |
| | | Theta | 0.71 ± 0.13 | 0.00 | 1.00 |
| Lee 2019 session 1 | 0.71 ± 0.12 | Alpha | 0.69 ± 0.10 | 0.02 | 0.09 |
| | | Beta | 0.70 ± 0.10 | 0.01 | 0.55 |
| | | Delta | 0.70 ± 0.10 | 0.01 | 1.00 |
| | | Gamma | 0.71 ± 0.11 | 0.00 | 1.00 |
| | | Theta | 0.70 ± 0.10 | 0.01 | 1.00 |
| Lee 2019 session 2 | 0.71 ± 0.12 | Alpha | 0.69 ± 0.11 | 0.02 | 0.12 |
| | | Beta | 0.70 ± 0.10 | 0.01 | 1.00 |
| | | Delta | 0.70 ± 0.11 | 0.01 | 0.90 |
| | | Gamma | 0.71 ± 0.11 | 0.00 | 1.00 |
| | | Theta | 0.71 ± 0.11 | 0.00 | 1.00 |
